# Recurrent Neural Networks with Interpretable Cells Predict and Classify Worm Behaviour

**DOI:** 10.1101/222208

**Authors:** Kezhi Li, Avelino Javer, Eric E. Keaveny, Andre E.X. Brown

## Abstract

An important goal in behaviour analytics is to connect disease state or genome variation with observable differences in behaviour. Despite advances in sensor technology and imaging, informative behaviour quantification remains challenging. The nematode worm *C. elegans* provides a unique opportunity to test analysis approaches because of its small size, compact nervous system, and the availability of large databases of videos of freely behaving animals with known genetic differences. Despite its relative simplicity, there are still no reports of generative models that can capture essential differences between even well-described mutant strains. Here we show that a multilayer recurrent neural network (RNN) can produce diverse behaviours that are difficult to distinguish from real worms’ behaviour and that some of the artificial neurons in the RNN are interpretable and correlate with observable features such as body curvature, speed, and reversals. Although the RNN is not trained to perform classification, we find that artificial neuron responses provide features that perform well in worm strain classification.

## 1 Introduction

As the principal output of the nervous system, behaviour plays a fundamental role in neuroscience and is also an important diagnostic criterion in psychiatric and motor diseases [1]. Quantifying behaviour is therefore of both scientific and technological interest. However, behaviour arguably encompasses the complete set of actions made by an animal over the course of its life, making it difficult to determine which aspects of behaviour to focus on. There is therefore growing interest in alternative representations and unsupervised methods to identify behaviours [2, 3, 4].

The nematode worm C. *elegans* has several useful features for testing new approaches for behaviour quantification. Its morphology is simple and it maintains a relatively constant length and width so that its motion can be captured by the coordinates of its midline over time and there are data available for thousands of individuals from several hundred genetically different strains [5]. Furthermore, the posture space defined by the angles of the midline is low-dimensional and can be well-described using a relatively small number of basis shapes [6]. This reduces the problem of behaviour analysis to finding patterns, features, or models that best describe trajectories through this shape space. Finally, because C. *elegans* is transparent and has a compact nervous system with only 302 neurons with well-described connectivity [7], mapping between neural activity and behaviour is more tractable than for many other animals.

In this paper, we test the hypothesis that a generative model that is able to produce realistic trajectories will need to learn essential aspects of the behavioural data it is trained on and therefore may also learn to distinguish animals based on their motion. To do this, we trained a Long Short-Term Memory (LSTM) recurrent neural network (RNN) to predict the next worm shape given previous shapes using back propagation through time. The trained model generates convincing behaviour that is difficult to distinguish from real behaviour and models trained on different strains generate trajectories that capture some of the known idiosyncrasies of those strains. We analyse the properties of the trained model and find cells with interpretable activation, including a cell related to reversals that is qualitatively similar to the activity of a real worm neuron measured in freely behaving animals. Notably, although the model is not specifically designed for classification, we achieve good performance on worm strain classification using artificial neuron derived features.

## 2 Model Training and Motion Prediction

We used a previously published data set consisting of 15 minute videos of single worms crawling on the surface of an agar plate [5, 2].

To describe the shape of the worm in each frame, we use the tangent angles of segments along the worm midline which are projected onto a lower-dimensional set of basis shapes, or ‘eigenworms’ [6]. Rather than using principal component analysis (PCA) [6] to find the basis shapes, we used independent component analysis (ICA) since it more effectively separates oscillating components [8]. We keep the top 6 shapes in our representation, which capture 98% of the variance across the mutant shapes, to ensure our model is able to capture even subtle motions that worms perform at the tip of their heads. The data that we use for training consists of the projected amplitudes 1 – 6 as well as the mean angle that was subtracted before projection to keep track of orientation (Fig. 1 and Supplementary Material).

**Figure 1:**
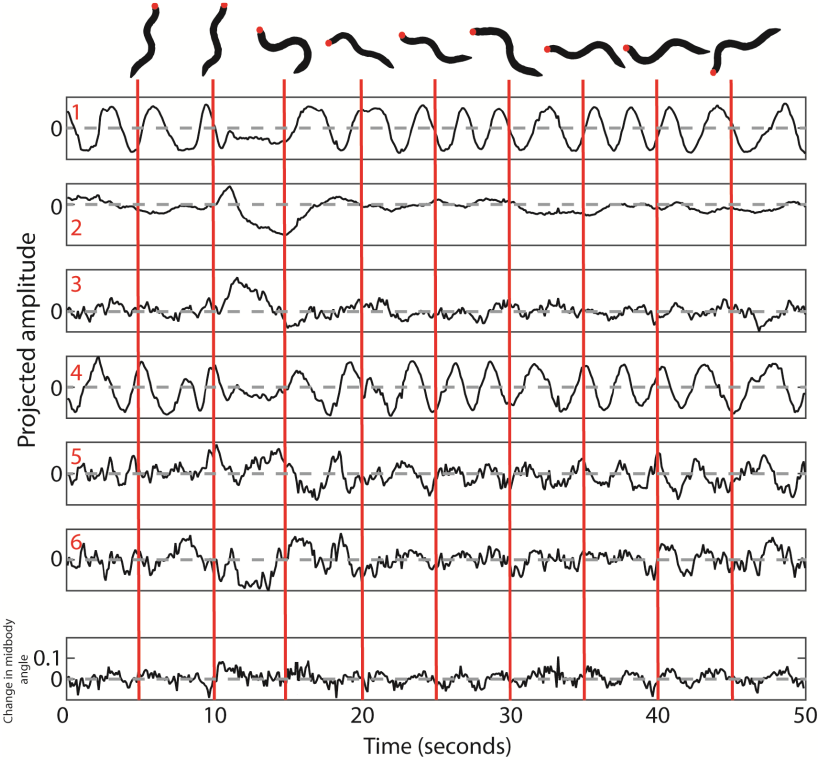
Projecting worm shapes onto the 6dimensional basis formed by the eigenworms plus 1-dimensional angle mean changes so a sequence of behavior can be compactly represented as a 7-channel time series.

We used Keras [9] to train an RNN with 4 hidden layers, each consisting of 260 LSTM [10] cells followed by a dropout cell to avoid overfitting [11, 12]. We chose to use LSTM cells because of their high performance in other domains. Using fewer than 260 cells per layer led to poor generative models (See Supp. Mat.) The last layer is a fully connected layer with a 7-dimensional vector as output. The data are subsampled to 10 frames per second and we used a 50 frame window to predict the next frame during training. The 5-second window used for training is on the same order as the body oscillation period of the worm.

We trained a separate model for each of four genetically different worm strains: the lab strain N2, two wild strains isolated from different parts of the world (CB4856, and ED3049), and a mutant *trp-4*(*sy695*). For the classification problem, we trained a single model on input data from 18 wild-isolates. Each model was trained using data from ten individuals (around 9 × 10^4^ frames in total per strain). The dropout rate was set to 0.2.

Each strain-specific model was seeded with 50 frames of real data and then used to generate 9000 frames of simulated data. After the initial seed, all subsequent frames are predicted by the model. We then used an updated linear resistive force theory algorithm to predict the worm’s *x* − *y* motion based on the shape changes [13] so that we could generate trajectories from the shape time series and extract motion features (such as speed) from the model-generated behaviour.

## 3 Results

### 3.1 Model generates realistic worm behaviour

Sample trajectories generated by the model are visually similar to those of real worms (Fig. 2). The model produces stable worm-like output even when extrapolating far beyond the size of the initial data seed. Since the *x* − *y* motion is predicted using a friction model from the shape changes and not as a direct output of the LSTM, the fact that the simulated trajectories look reasonable is evidence that the model has learned essential aspects of worm behaviour. Consistent with applications of LSTMs in text analysis [14], we find that the simulated trajectories have realistic behaviour but at time scales longer than the 50-frame training window including active ‘roaming’ periods, inactive ‘dwelling’ periods, and sharp turns and reversals. The simulated trajectories also show more gradual turning during forward locomotion, again consistent with the behaviour of real worms.

**Figure 2:**
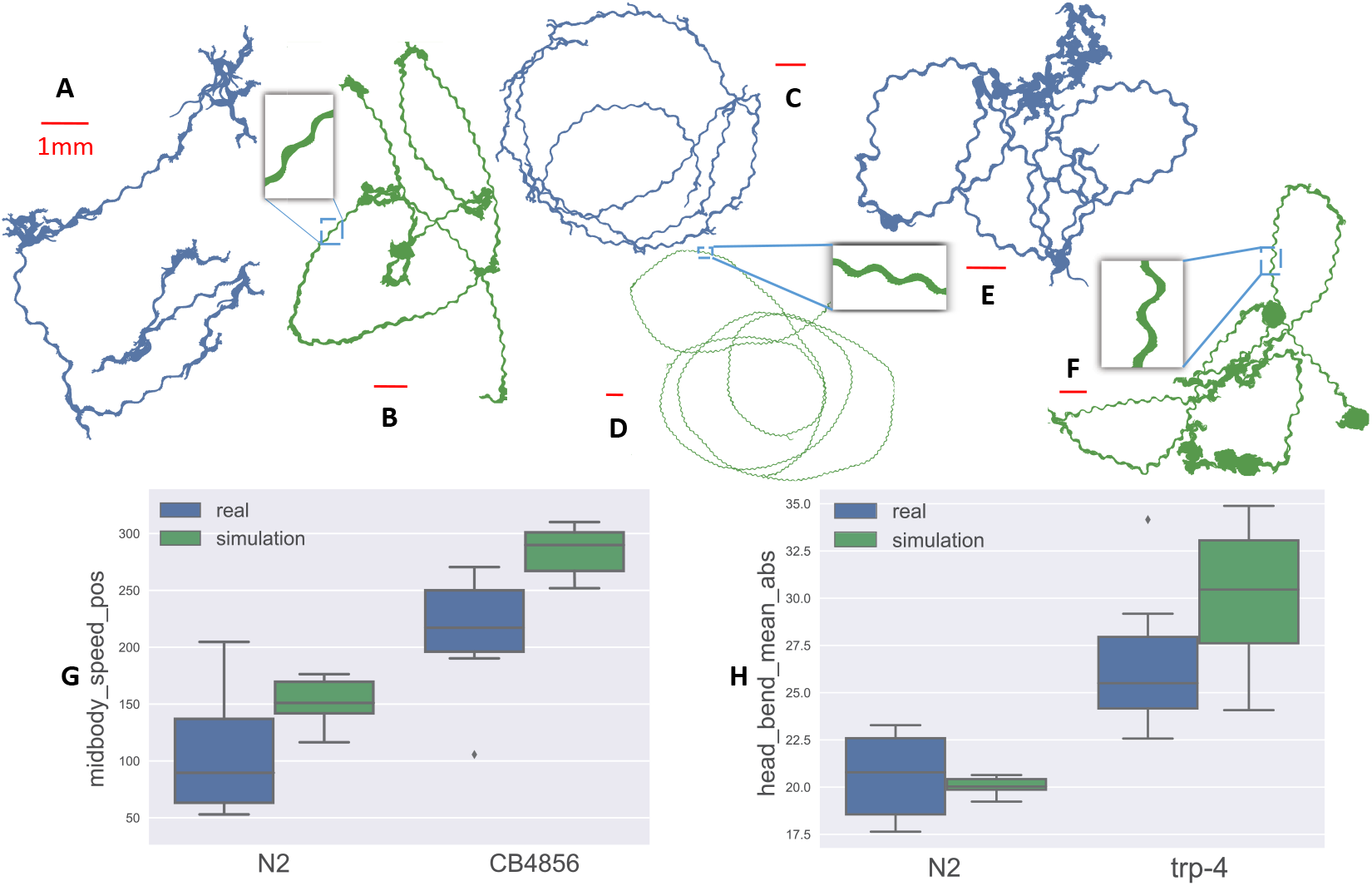
Real and simulated trajectories of different strains. A: real N2; B: simulated N2; C: real CB4856; D: simulated CB4856; E: real *trp-4*; F: simulated *trp-4*; G: the boxplot of real vs. simulated worms in terms of the feature *midbody_speed_pos* for N2, CB4856; H: the boxplot of real vs. simulated worms in terms of the feature *head_bend_mean_abs* for N2 and *trp-4*.

Animals with deletions of the gene *trp-4* have a more curved body posture because of a defect in proprioception [15] and CB4856, a wild strain isolated from Hawaii, is more active than the laboratory strain [16]. We generated simulated trajectories from models trained separately on data from each of these strains. The expected differences are visible in the real and simulated trajectories shown in Figure 2. These differences were consistent across 30 simulated trajectories using different seeds for each run (Fig. 2,G and H). However, we also note the match is not perfect, confirming that the model is not simply returning shape sequences already present in the training data, which might have been expected if the models were overfit.

### 3.2 Interpretable LSTM cells and interpretable worm cells

Recent work has shown that interpretable cells, that is cells whose memory correlates with an observable property of the modelled system, can exist in LSTMs trained on real-world data [14, 17]. When modelling behaviour, where we know that the training data were themselves generated by a neural network, it is plausible that there could be an informative mapping between the artificial and real neurons. In the case of *C. elegans,* this mapping may be more tractable than in other systems because the worm has only 302 neurons with a known wiring diagram [7] and because it lacks voltage gated sodium channels and therefore has non-spiking neurons [18].

By comparing the variation of LSTM cells’ memories over time with the features we routinely calculate for worm behaviour, we identified cells with interpretable behaviour. In the 4th layer, we find several cells with a strong correlation or anti-correlation (correlation coefficients above 0.8 or below −0.8) with the worm’s midbody curvature. Given that some of the basis shapes used in our representation encode curvature fairly directly, the presence of such cells is not surprising. However, we also discovered single units that anti-correlate with speed (Fig. 3), a feature that is not directly present in the input shape data (correlation coefficient −0.42). The relationship is clear upon visual inspection when both signals are smoothed with a 200-frame mean filter (correlation coefficient −0.75 after smoothing).

**Figure 3:**
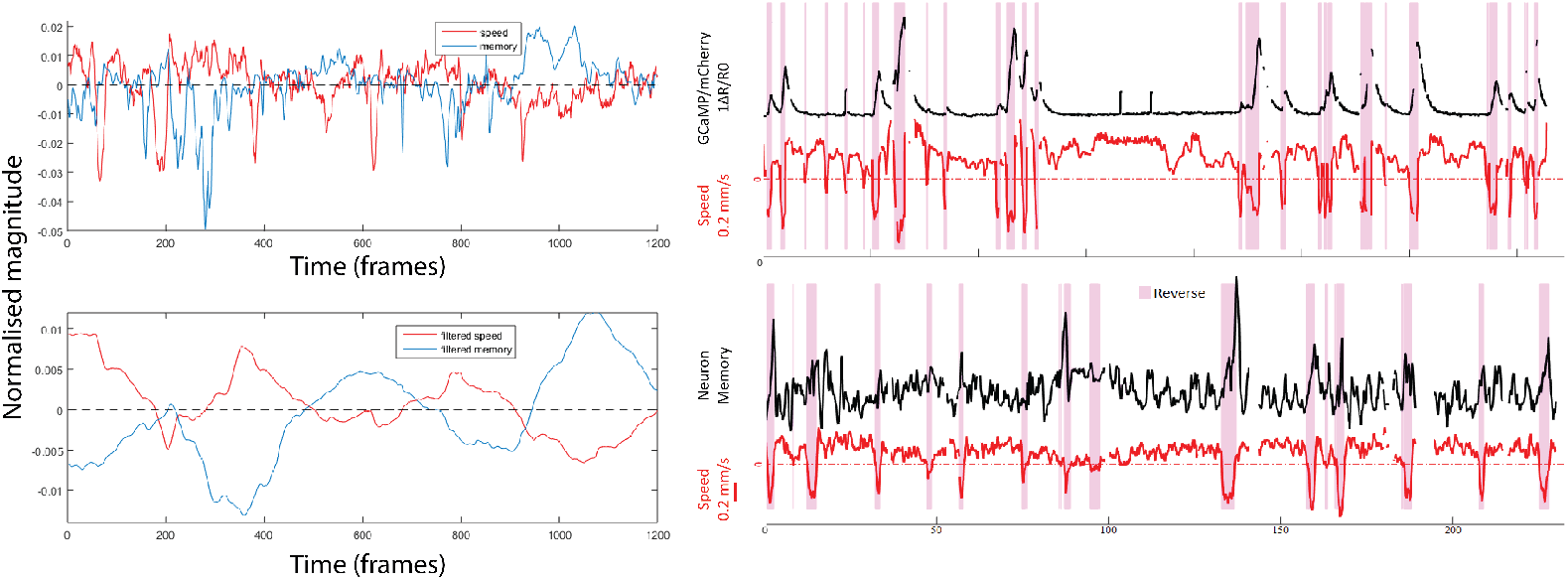
Left: Time series of midbody speed (red) vs. LSTM memory (blue) (upper) and after a 200 frame mean filter (bottom). Right: Sample trace showing the activity of RIM, a real worm neuron (data from [19]), measured using a fluorescent *calcium* reporter in a freely behaving animal (fluorescence ratio (black) and corresponding crawling speed (red)). Time series of LSTM memory (black) and midbody speed (red) (bottom).

Finally, we found an LSTM cell that is more active when the worm is moving backwards (Fig. 3). Interestingly, the RIM neuron in the real worm has a similar relationship to reversals, as shown by the calcium (Ca^2+^) imaging dynamics in 3 (data from [19]). We do not find neurons that correlate with, for example, dwelling states or omega turns, so we do not think the relationship between this LSTM cell’s activation and reversals is a result of having enough neurons to find arbitrary correlations by chance.

### 3.3 The generative model is also a good classifier

Some artificial neurons correlate with manually defined features, but not all do, suggesting the neural features may contain independent information. We therefore checked whether the artificial neuron activities could themselves be useful features for distinguishing strains. We trained a single RNN on data from 18 wild isolate strains of *C. elegans* and then fed data from individuals not included in the training set to the network and recorded the response of each artificial neuron. We used the 10th, 50th, and 90th percentile values of each neuron to generate a feature vector for each of the unseen worms, followed by recursive feature elimination to find the top 30 features. Simple nearest neighbour classification using the neural features achieved 52% accuracy using 5-fold cross-validation. This outperforms the classification using the manually defined features from Yemini *et al.* [5] which achieved 19% accuracy using nearest neighbours, and is comparable to the 56% (also 5-fold crossvalidation) achieved using a random forest with the Yemini *et al.* features. Given that the Yemini *et al.* features have previously been shown to sensitively distinguish worm strains from each other [5] and that the RNN was not trained on classification, this result was unexpected.

## 4 Conclusion

Quantifying behavioural dynamics remains challenging and there are still no general frameworks that are effective for all problems. Following training, LSTMs are able to generate realistic trajectories that capture known behavioural differences between genetically distinct worm strains. Although trained on prediction, we also found that the LSTMs could achieve state-of-the-art performance on strain classification.

We also found that the LSTMs have several interpretable cells. In at least one case the artificial neuron’s activity could be direclty related to the activity of real worm neurons. Of course, the LSTM used to model worm behaviour is very different from the worm’s actual nervous system, not least because it takes in sensory input and outputs patterns of muscle contraction whereas the LSTM has posture time series as both input and output. Nonetheless, many of the real worm neurons are not currently ‘interpretable’. Finding LSTM cells that correlate with the activity of these enigmatic real neurons could provide a complementary approach to understanding their role using the LSTM as a more readily perturbed system to study these cells’ effects on behaviour.

## Supplemental Material

### 1 The Model

We use a previously published data set that was captured using a USB microscope mounted on a motorised stage [1,2]. Each individual animal was recorded for 15 minutes at approximately 30 frames per second using the Worm Tracker 2.0 (WT2) system (http://wormbehavior.mrc-lmb.cam.ac.uk/). In the videos, worms are confined to the two-dimensional surface of an agar plate. We have updated the tracking and stage alignment algorithm from [1] to improve the speed and accuracy.

The deep learning development toolkit Theano (http://deeplearning.net/software/theano/) is used to implemented the RNN. We adopted the Keras Sequential model for training [3]. Our model has 4 LSTM layers, each followed by a dropout layer. The output layer is a fully connect dense layer. For the LSTM layer, each layer has 260 units. The dropout is set as 0.2. The input data are 7-dimension time series after dimension reduction of worm posture. The loss function is mean squared error, and the optimizer is set as the rmsprop optimizer in Keras. No regularizer is adopted in the model and the activation function of the dense layer is linear. It takes 600 epochs (each epoch scans through all the training frames exactly once) to obtain the model for one strain.

### 2 Dimension reduction

We use the skeleton, or midline, to describe the shape of the worm at each frame k, defined as a curve that passes through the centre of the body (Fig.1.A in red). The skeleton consists of a series of points *s*(1),…, *s*(*n*), where n is the number of points on the skeleton from head to tail and *θ*(*s*(*i*)) is the angle between the horizontal line and a vector connecting two consecutive points *s*(*i* + 1) and *s*(*i*) (Fig.1.B). By subtracting the mean angle, equivalent to rotating the worm shape by an angle *s*′ (Fig.1.C, solid line) we achieve a position and orientation independent representation of the worm shape in each frame [4]. Given worm shapes across many videos, we can calculate a set of basis shapes *u_μ_*(*s*),*μ* = 1,…*μ′* that can be used to decompose *θ*(*s*) to μ′ dimensions as a superposition of ‘eigenworms’

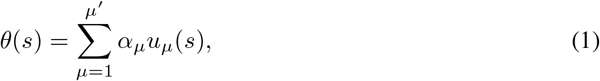

where*α_μ_* are the amplitudes of the projections onto the eigenworms. Rather than using principal component analysis (PCA) [4] to find the basis shapes, we used independent component analysis (ICA) since it more effectively separates oscillating components [5]. We keep the top *μ′* = 6 shapes in our representation, which capture 98% of the variance across the mutant shapes (Fig.1.D). The basis shapes over all mutants are shown in Fig.I.E.

**Figure 1:**
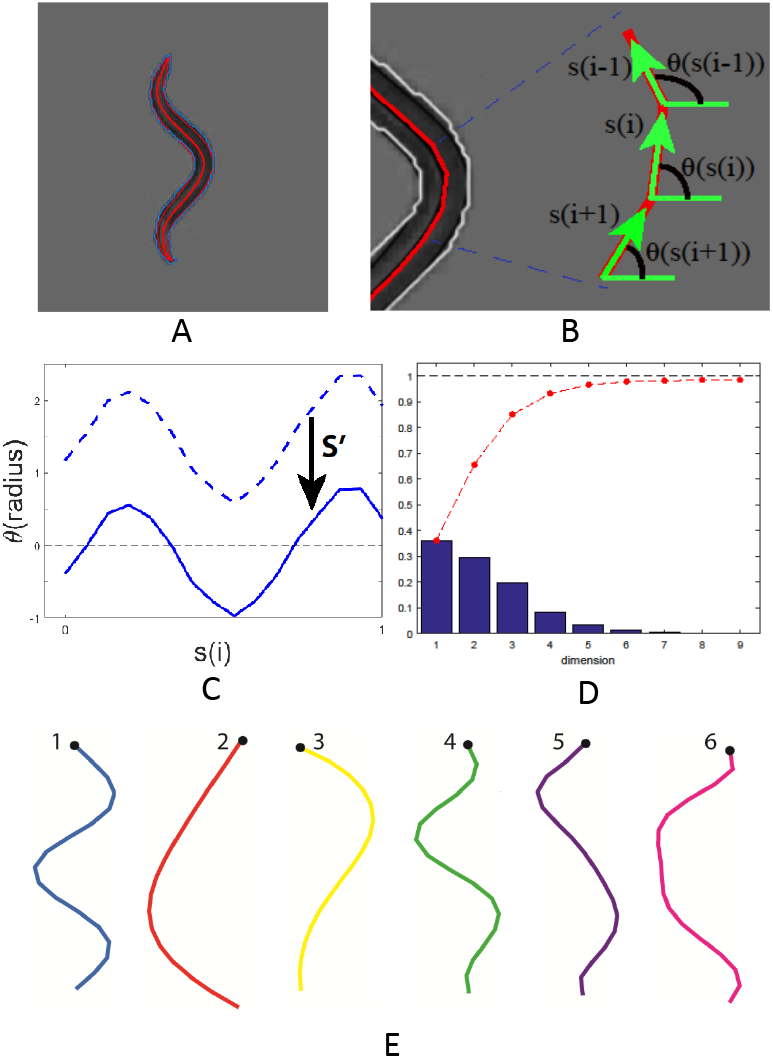
A: an example of a typical worm video frame with the skeleton identified. B: the definition of points s and angles *θ*. C: the curve of angles and the adjustment by its mean value s′. D: fraction of variance explained as more basis shapes are used. E: ICA returns 6 basis shapes that explain 98% of the variance in the dataset.

## 3 Model Training and Motion Prediction

### 3.1 Training

Because of their successful applications in modelling sequential data [6], we used a multi-layer LSTM [7] RNN to model the worm behaviour time series. We adopted the Keras Sequential model for the training component [3]. In general, sequence learning tries to map an input sequence to a target sequence of fixed-size. An LSTM RNN estimates the conditional probability *p*(*y*_1_,…, *y_T′_*|*x*_1_,…, *x_T_*) given a sequence of data, where *x*_1_,…, *x_T_* denotes the input sequence and *y*_1_,…, *y_T′_* is the corresponding output sequence, with size *T* and *T′* respectively. We use *x_t_*, *h_t_*, *c_t_* to represent input vector, output vector and memory cell vector respectively; *W*, *U*, and b are the parameter matrices/vectors that need to be learned, and *f_t_*, *i_t_*, and *o_t_* denote the forget gate, input gate, and output gate vectors. Then the precise form of the update is

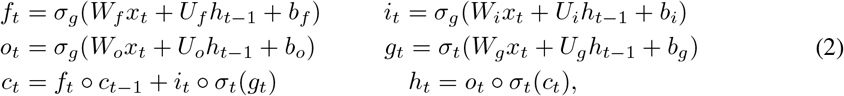

where*σ_g_*, *σ_t_*, o is the sigmoid function, hyperbolic tangent and entrywise product, respectively. A schematic of the data processing pipeline is shown in Fig. 2. The worm video is segmented and the skeleton is converted to a 7-dimensional time series that is used to train the RNN. We feed the last 50 time points to the model and use the next-frame values as outputs, stepping through the data one frame at a time. The data are subsampled to 10 frames per second and so 50 frames is of the same order as the period of worm body oscillation.

**Figure 2:**
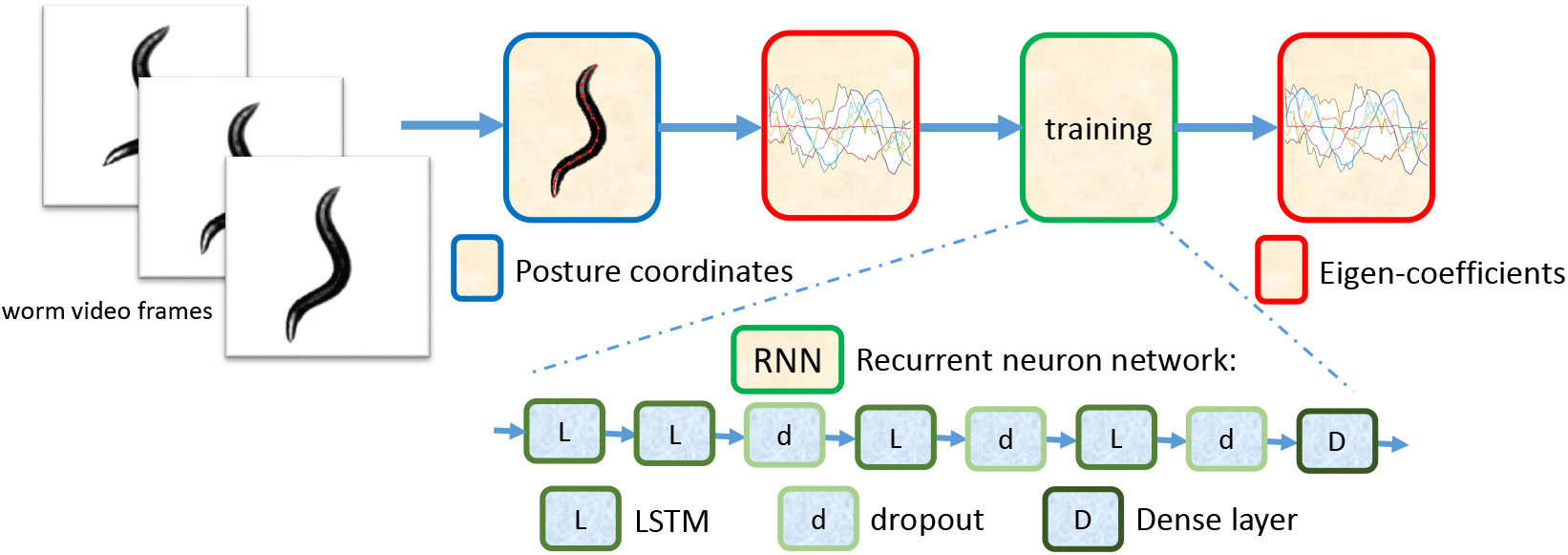
Worm behaviour is represented as a low dimensional time series. The posture coordinates are the *x* − *y* coordinates of the points on the worm’s skeletons. The input and output of the training component are time series data after dimensionality reduction. The LSTM has 4 hidden layers, each followed by a dropout cell, with a fully connected final layer.

**Figure 3:**
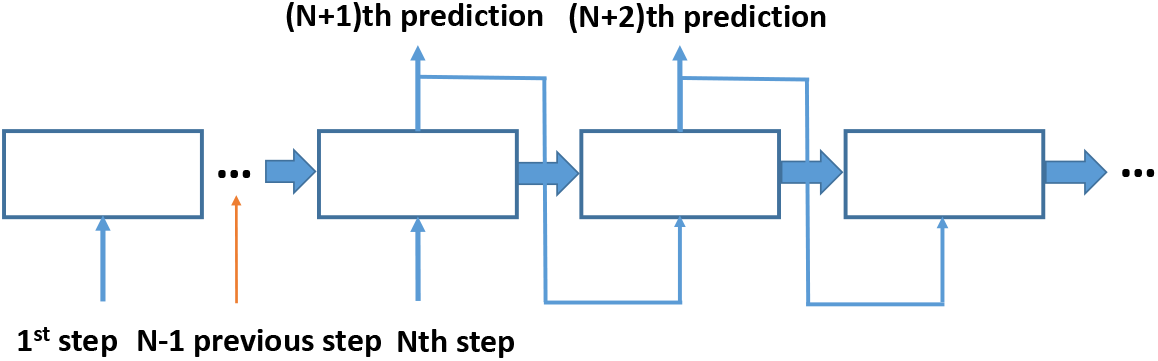
Schematic of prediction algorithm using an LSTM.

### 3.1.1 Model Size Investigation

In the paper we trained an RNN model with 4 hidden layers, each consisting of 260 LSTM cells. We tried smaller networks but did not perform as well as generative models. Sample trajectories are shown in Fig. 4.

**Figure 4:**
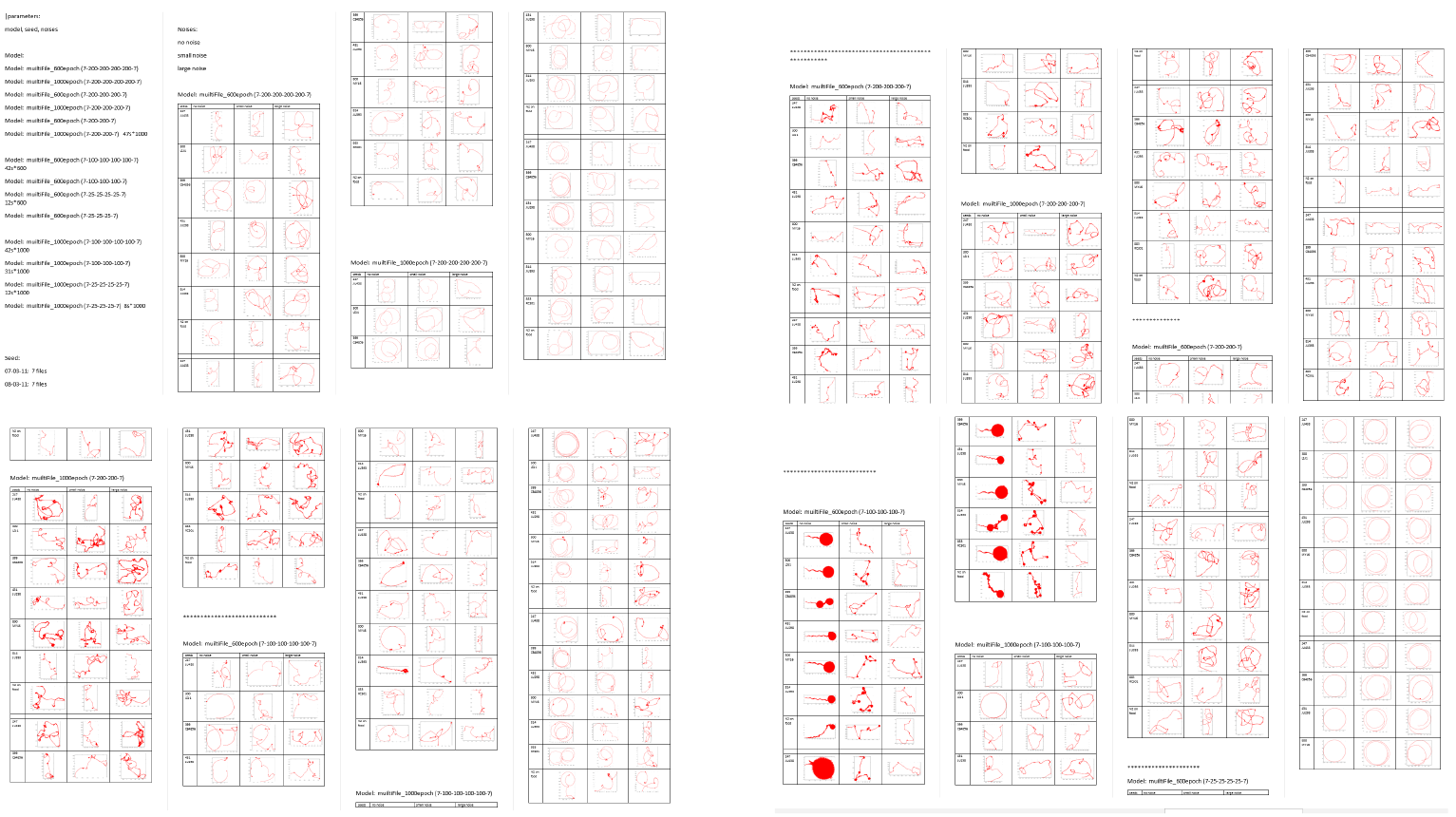
Sample trajectories generated with RNNs with smaller numbers of neurons than the one used in the main text.

**Figure 5:**
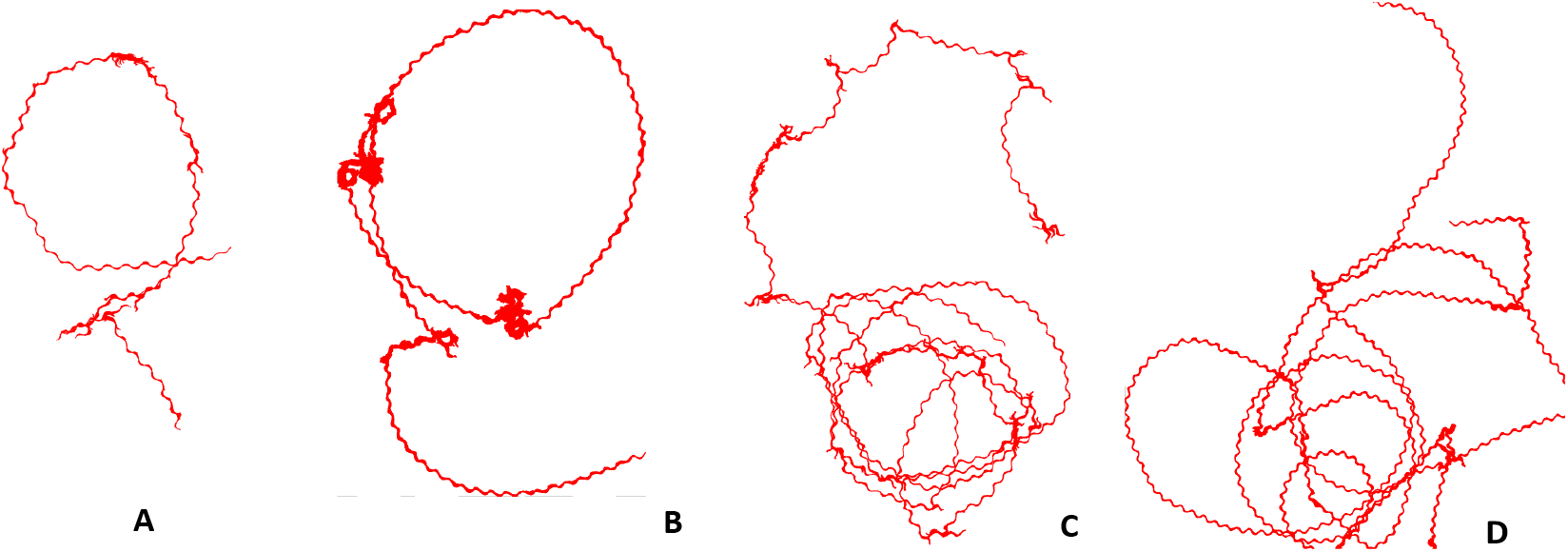
Trajectory examples; A: real N2, B: simulated N2, C: real ED3049, D: simulated ED3049

### 3.1.2 Real-or-Simulated test

Worm behaviour is more complex and diverse than the trajectories can illustrate because they do not show differences in dynamics very clearly. To further assess the apparent similarity between the real and simulated behaviour, we generated short 1-minute videos of real worm behaviour and simulated worm behaviour, in each case displaying only the skeleton coordinates over time, and asked volunteers to classify the clips as real or simulated. We divided the volunteers into two groups: researchers working with *C. elegans* and those with no prior experience with *C. elegans*. After watching 5 to 10 labelled examples to familiarise them with the task, users were given unlabelled videos to watch and could press a button indicating that they thought the video was real or simulated at any time during the video (Fig. 6). Both real and simulated videos only show the skeleton (in red) and the head point (in blue) of the worm.

**Figure 6:**
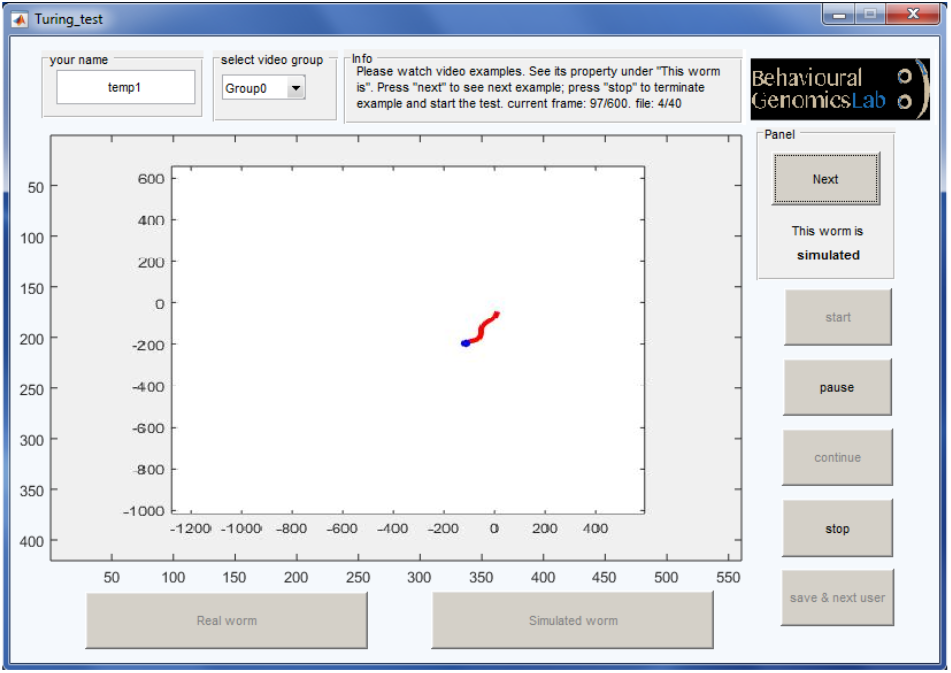
The programme interface shows examples of *real* or *simulated* videos before the formal test.

**Figure 7:**
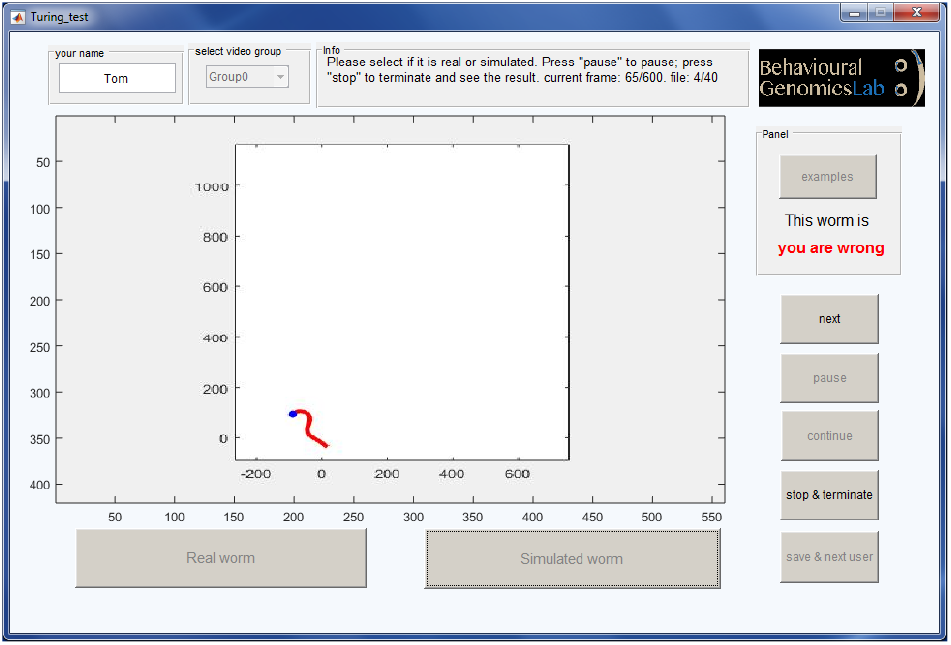
The programme interface shows the correct identity immediately after an answer is submitted during the formal test.

The correct answer will show immediately after the user makes a choice, and then the program goes to the next video (Fig. 7). The test results are shown in Table 1.

**Table 1:**
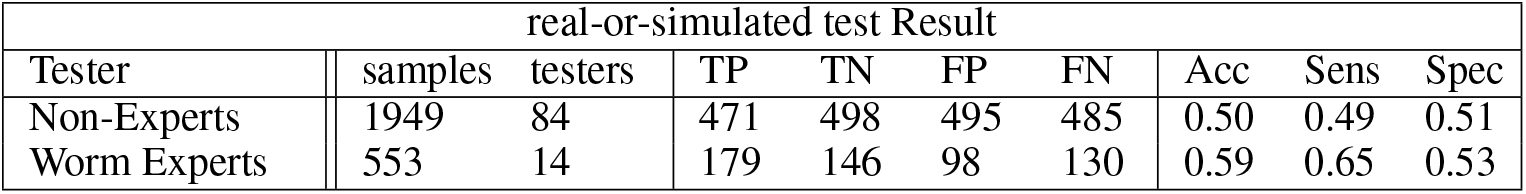
Test results of real-or-simulated test to distinguish real or simulated worm videos. The abbreviations in the table stand for: true positive, true negative, false positive, false negative, accuracy, sensitivity and specificity, respectively.

| real-or-simulated test Result |  |  |  |  |  |  |  |  |  |
| --- | --- | --- | --- | --- | --- | --- | --- | --- | --- |
| Tester | samples | testers | TP | TN | FP | FN | Acc | Sens | Spec |
| Non-Experts | 1949 | 84 | 471 | 498 | 495 | 485 | 0.50 | 0.49 | 0.51 |
| Worm Experts | 553 | 14 | 179 | 146 | 98 | 130 | 0.59 | 0.65 | 0.53 |

In the table one can observe that non-experts achieved a test accuracy around 50%, equivalent to random guessing. The results show that even people working with *C. elegans* on a daily basis have difficulty distinguishing the real from simulated worms.

## 4 Interpreting Cells

For visual inspection, we check units during a 1-frame prediction by visualizing their values as the worm video plays in realtime (Fig. 8). The interface shows the worm shape and location in the video (upper left), the specific unit which is under observation (upper right), and the realtime unit values in each LSTM layer. We also can show the unit values in the format of time series data, which have been shown in Sec. Interpreting LSTM Cells in terms of *bend, speed* and *reversal*.

**Figure 8:**
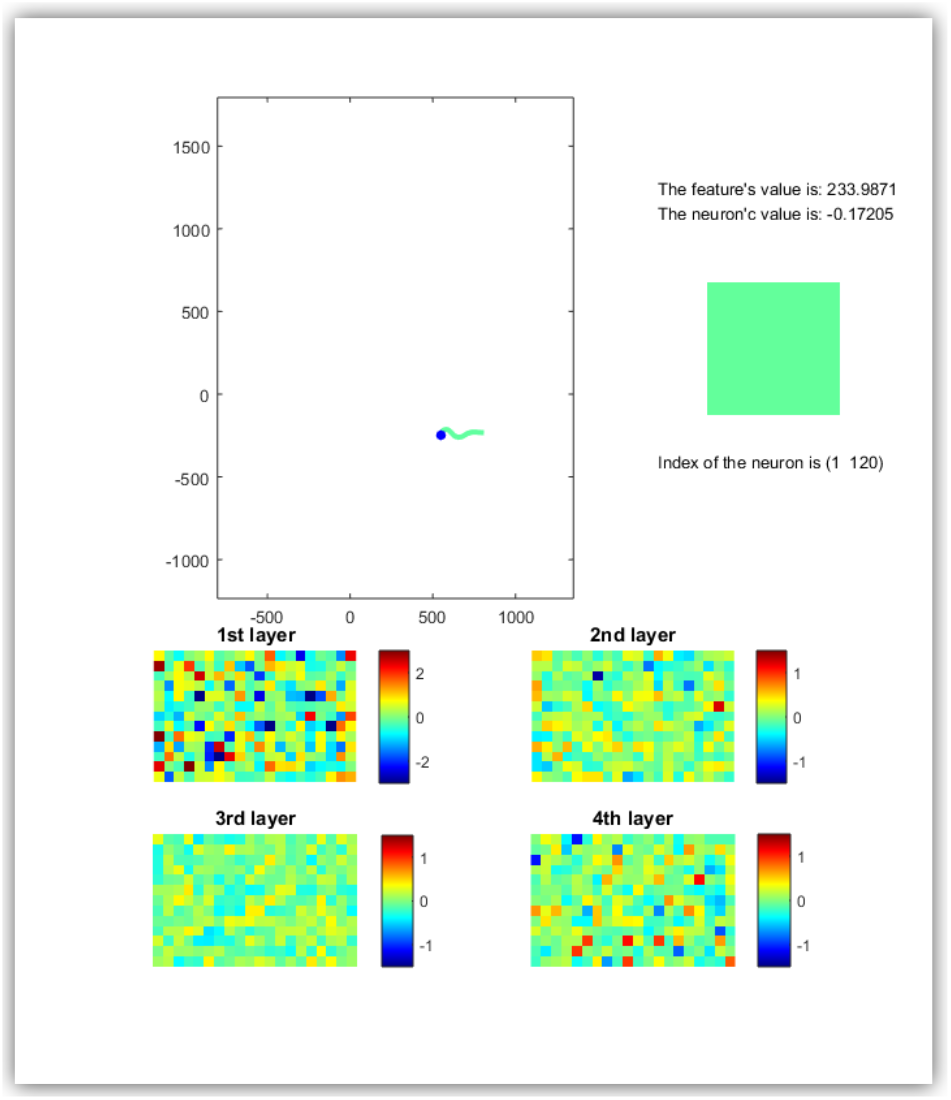
The interface showing the values of units in each LSTM layer during the worm video proceeds realtime.

## Acknowledgments

We thank Bertalan Gyenes for help drawing Figure 1, Linus Schumacher, Serena Ding and Angie Yin and the Imperial Festival for help with the Real-or-Simulated tests.

